# Local rabies transmission and regional spatial coupling in European foxes

**DOI:** 10.1101/710988

**Authors:** Laurie Baker, Jason Mathiopoulos, Thomas Müller, Conrad Freuling, Katie Hampson

## Abstract

Infectious diseases are often transmitted through local interactions. Yet, both surveillance and control measures are implemented within administrative units. Capturing local transmission processes and spatial coupling between regions from aggregate level data is therefore a technical challenge that can shed light on both theoretical questions and practical decisions.

Fox rabies has been eliminated from much of Europe through oral rabies vaccination (ORV) programmes. The European Union (EU) co-finances ORV to maintain rabies freedom in EU member and border states via a cordon sanitaire. Models to capture local transmission dynamics and spatial coupling have immediate application to the planning of these ORV campaigns and to other parts of the world considering oral vaccination.

We fitted a hierarchical Bayesian state-space model to data on three decades of fox rabies cases and ORV campaigns from Eastern Germany. Specifically, we find that (i) combining regional spatial coupling and heterogeneous local transmission allows us to capture regional rabies dynamics; (ii) incursions from other regions account for less than 1% of cases, but allow for re-emergence of disease; (iii) herd immunity achieved through bi-annual vaccination campaigns is short-lived due to population turnover. Together, these findings highlight the need for regular and sustained vaccination efforts and our modelling approach can be used to provide strategic guidance for ORV delivery. Moreover, we show that biological understanding can be gained from inference from partially observed data on wildlife disease.

## Introduction

Disease dynamics are underpinned by the interplay between population connectivity and the localized nature of transmission [1–4]. Many infectious diseases are transmitted primarily through local interactions, but control strategies and surveillance are implemented at coarser administrative scales. As a result, only aggregate data is available on the occurrence of infections, which makes it difficult to disentangle the extent to which disease dynamics are driven by local transmission versus spatial coupling between subpopulations. Approaches that quantify these processes have potential to guide the efficient use of resources for disease control.

Epidemiological data are inherently complex due to variability arising from infectious disease dynamics, including stochastic spatial transmission processes and observation errors in detecting cases. The study of wildlife disease is particularly complex for several reasons. Surveillance only detects a (usually small) proportion of circulating infections and oftentimes the incidence of disease in wildlife populations can only be inferred through indirect measures (veterinary records, hunting reports), with variable levels of detection [5]. Under limited surveillance failed invasions or low level persistence may be missed entirely. Wildlife populations are often not monitored closely and therefore knowledge of their size and spatial distribution are typically limited and imprecise [6]. Animals and humans move in different ways, as a result of social groups, territorial boundaries, and specific habitats and geographical features that can direct or impede movement [7, 8]. Many animals have faster demographic rates than humans [9], such that the herd immunity achieved through vaccination is relatively short-lived. Finally, vaccination programmes targeting wildlife are challenging to implement and monitor [10–12].

Traditional epidemiological models that assume homogenous mixing i.e. individuals interacting randomly and uniformly with all others in the population, have yielded important insights, such as thresholds for disease invasion and control [13–15]. But these models have not accounted for heterogeneous mixing, which is critical for directly transmitted diseases in wildlife populations. Individual-based models explicitly model interactions within discrete spatial or social neighbourhoods [1, 16], but require detailed data that is rarely available for wildlife populations. Approximations have been developed to capture interactions between infected and susceptible individuals at the local level [17–21], for example, ‘heterogeneity’ parameters [17, 18, 22]. These approaches have been effective for human diseases such as cholera [23] and measles [1], but have not been applied to wildlife diseases such as rabies.

Rabies has been eliminated from fox populations throughout much of Europe by vaccinating foxes using oral baits containing vaccine. In just over three decades, vaccine baits have been distributed across 2.36 million km^2^ [24–26]. Since the late 1980s the European Union (EU) has co-financed Oral Rabies Vaccination (ORV) programmes in member and border states [24, 25, 27]. Models that capture local transmission dynamics of fox rabies and regional connectivity therefore have immediate application to the situation in Europe and elsewhere. Here, we examine fox rabies dynamics in response to oral vaccination using a hierarchical Bayesian state-space model fit to incidence data from Eastern Germany from 1982-2013. We use a metapopulation approach to model transmission by representing space as a network of subpopulations and estimating the movement of infected individuals (or coupling) between them. We account for heterogeneous mixing using a transmission process that approximates the scaling of individual interactions to the regional level. This study presents a first step towards disentangling local transmission and spatial coupling between subpopulations from aggregated and incomplete data on wildlife rabies.

## Materials and Methods

We analysed monthly time series of fox rabies cases for the period 1982-2013 from 5 federal states in Eastern Germany (Brandenburg, Mecklenburg-Vorpommern, Sachsen, Sachsen-Anhalt, and Thüringen) in relation to the timing of ORV campaigns, fitting a hierarchical Bayesian state-space model to these data.

### 0.1 Data Collection

We compiled records of laboratory-confirmed rabies cases in foxes from regular reports by the national veterinary authorities and summarized for each federal state (hereafter referred to as region) on a monthly basis. Specimens of suspect rabid foxes were submitted primarily by veterinarians and hunters. From 1993, cross-sectional sampling of foxes was also conducted, whereby a proportion of foxes hunted were tested for rabies providing a measure of rabies prevalence in the population. The timing of ORV campaigns in each region was also compiled. The Rabies Bulletin Europe (RBE) is a database consisting of national rabies surveillance data managed by the WHO Collaborating Centre for Rabies Surveillance and Research at the Friedrich-Loeffler-Institut in Germany [28, 29]. A monthly average number of confirmed rabies cases was calculated from the RBE quarterly reports for neighbouring regions in Poland and the Czech Republic that border the five federal states in Germany.

### 0.2 Bayesian State-Space Model

A discrete time stochastic metapopulation model with three states: Susceptible (S), Infected (I), and Vaccinated (V), was developed to model the numbers of foxes and rabies cases in different regions through time. The Bayesian approach allowed us to complement the rabies case data with prior information on some parameters from historical studies on fox demography.

A demographic process was used to model the numbers of susceptible and vaccinated foxes in each region at monthly time steps. The starting susceptible population and carrying capacity for each region were extracted from the literature, based on the average density of foxes per km^2^ scaled up to the region [30–32]. Births were modelled as occurring in April of each year with newborn foxes entering the susceptible population in July coinciding with when they venture further from their den. All susceptible and vaccinated foxes older than one year of age were considered reproductively active. This means that surviving newborns from the past year give birth the following April. We assume that infected individuals transmit rabies and die within the same month, such that infectious animals in the current month *t*, transmit infection to new animals that subsequently develop rabies the following month *t* + 1. No exposed class was considered because the latent period of rabies infection lasts an average of three weeks and thus all new infections at month *t* become symptomatic by month *t* + 1 [33]. Data on the timing of vaccination campaigns in each region were incorporated explicitly.

Susceptible individuals *S* in region *r*, in month *t* were modeled as a function of juvenile foxes entering the susceptible population three months after birth *j*_*r,t*_, surviving individuals *C*_*r,t*_ and those removed due to vaccination *V*_*r,t*_ or infection *I*_*r,t*_:

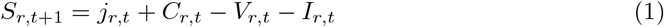

The first term *j*_*r,t*_ is a binomially distributed variable representing juvenile foxes entering the susceptible population three months after birth and takes the form:

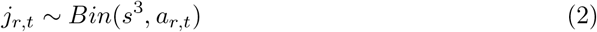

where *s*^3^ is the 3-month survival probability and *a*_*r,t*_ are the newborn foxes. Foxes live to a maximum age of about 4 years [32]. If we assume that only 1% of foxes are alive at age 4 years we can use the following expression to determine the survival probability: *s*^48^ = 0.01, *s* = 0.01^(1/48)^ = 0.909

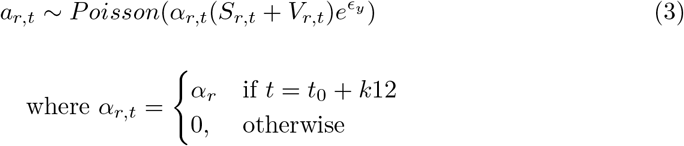

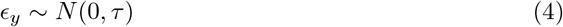

and *α*_*r*_ is the per capita annual birth rate in region *r* in month *t* applied to all susceptible *S* and vaccinated *V* individuals in the system. Fecundity is regulated by annual fluctuations in the environment, 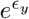. Here, we use the exponential term 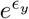 and a normal prior centred around 0 for *ε*_*y*_ to capture the effect of environmental noise on the size of the birth pulse. Under the exponential, when *ϵ* is 0, then *e*^0^ = 1, meaning there is no change in the size of the birth pulse. An *ε* smaller or greater than 0 will result in a smaller or larger birth pulse, respectively. The prior for the precision term, *τ*, is specified such that the birth pulse can vary by +/−10% in line with fluctuations in the birth pulse observed in wild fox populations [34].

The realised per capita annual birth rate with density dependence takes the form:

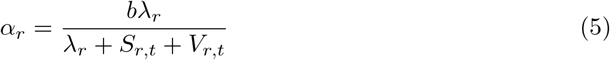

where *b* is the maximum annual per capita reproductive rate. Here, the inclusion of *S* and *V* leads to density dependence, the strength of which is controlled by the parameter *λ*_*r*_ (derived in Appendix **??**) in the different regions *r* and takes the form:

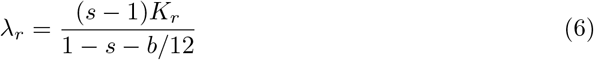

where *s* is the survival probability (see Eq (2)) and *K*_*r*_ is the carrying capacity in region *r*.

The second term in Eq 1 comes from a binomial distribution and represents the surviving susceptible individuals in region *r* and month *t*,

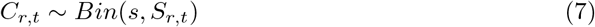

where *s* is the survival probability (see Eq (2)) and is assumed to be fixed across time and regions.

Infected individuals are modelled as:

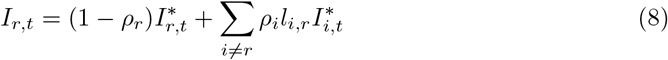

where 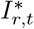 is the number of infected individuals in region *r* prior to any movement. Infected individuals leave the region with probability *ρ*_*r*_. The summation term represents incursions from other regions as a function of rabies incidence 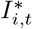 in region *i*, the proportion of infected animals leaving region *i*: *ρ*_*i*_, and *l*_*i,r*_ the proportion of those moving to region *r*. This can be modelled in different ways, but here we chose to equate it to the proportion of the border of region *i* that is shared with region *r*.

The probability of leaving *ρ*_*i*_ was calculated as:

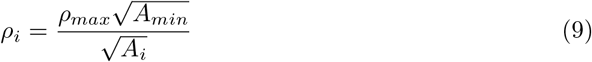

where *ρ_max_* is the maximum leaving rate. Under a diffusive assumption for movement the leaving rate is expected to decrease as the size of the region increases relative to its perimeter and the probability that any given infected animal moves outside of its region declines. To capture this effect, we scaled the leaving rate by dividing it by the area of the region 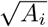, which is proportional to how far individuals are from the perimeter of the region.

In the model, new infections are generated within region *r* from a Binomial distribution:

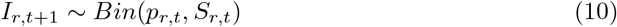

where the new infected individuals *I*_*r,t*_ at time *t* + 1 in region *r* are generated from the susceptible individuals *S* with a risk of transmission probability *p*_*r,t*_ represented by:

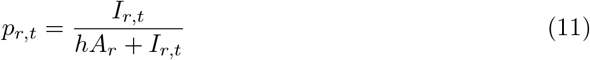

Here *hA*_*r*_ is the half-saturation point for *p*_*r,t*_. That is the number of infected individuals that raise the transmission probability to 0.5. We assume that this number is only a function of the total area of the region, appropriately scaled by the constant *h* which is estimated from the data. Subsequently, the risk of infection *p*_*r,t*_ depends on the density of infected individuals 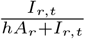in the region, with the transmission rate per infected individual decaying as the number of infected individuals grows.

The biological rationale behind Eq (11) is that in larger areas, susceptible individuals are expected to be less accessible to infected individuals due to the greater distance to reach them. The addition of infected individuals in the denominator allows us to account for saturation effects that occur at high incidence, when infected individuals might contact fewer susceptibles due to disease-induced mortality or because local contacts might already be latently infected. Local susceptible depletion thereby reduces the transmission rate of each infected individual as incidence increases.

The functional form of transmission is a special case of the general model in Eq 35 from [18]:

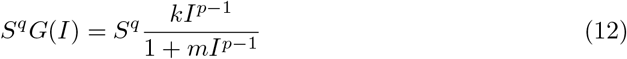

where *S* is the susceptible population, *I* is the number of infected individuals and k, p, q, and m are positive constants. If we assume that q is 1, G(I) can be written as

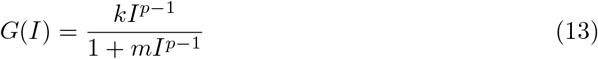

If we consider the special case where *p* = 2, *m* = (*hA*)^−^1, *k* = *hA* and *q* = 1 then

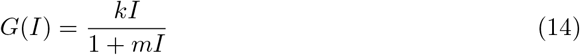

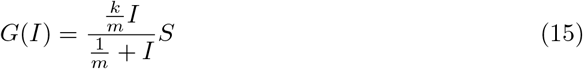

which is equivalent to Eq (11).

The time evolution of the number of vaccinated individuals is modelled as:

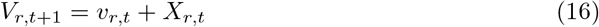

where *v*_*r,t*_ represents the newly vaccinated individuals drawn from a binomial distribution.

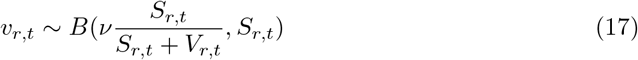

and *ν* is the rate of bait uptake by the population of susceptible and vaccinated individuals in region *r* at time *t*. The bait uptake rate *ν* was given a fixed Beta prior with mean 0.30 and variance of 0.005 based on field studies [35]. To account for the depletion of baits by already vaccinated conspecifics, the rate of bait uptake by susceptible individuals is determined relative to their proportion in the population 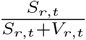. Vaccination is switched on and off by an indicator variable that is 0 in all months apart from those when a vaccination campaign occurred when it equals 1. Vaccination campaigns were typically carried out in September or October and April during the study period. The term *X*_*r,t*_ in Eq (16) represents surviving vaccinated individuals from the previous time step drawn from a Binomial distribution.

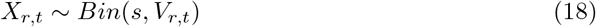

where *s* is the survival probability (see Eq (2)).

We assumed that infected individuals *I*_*r,t*_ were observed imperfectly each year with probability *θ*_*y*_:

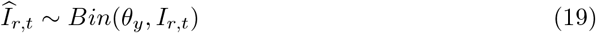

that varies stochastically on an annual basis.

In the cross-sectional sampling regime, hunted foxes *H*_*r,t*_ had a probability of being observed to be infected equal to the risk of transmission *p*_*r,t*_.

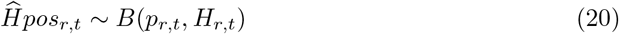

where 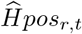 is the number of positive cases out of the total foxes hunted *H*_*r,t*_.

### 1 Model Fitting

All models were fitted using the software JAGS [36], which uses Gibbs sampling to generate posterior distributions of the parameters given the likelihood, prior distributions and the data itself. We ran the models for 3,000,000 iterations, with a burn-in of 30,000 and a thin interval of 300, giving 10,000 samples. We inspected the model for convergence and effective sample size. To account for the fact that fox-mediated rabies had been circulating in Germany since the late 1940s, we started the model 10 years (120 time steps) prior to when the time series began to allow the system to settle at an endemic equilibrium. Fitting of the model required considerable computational time and a compromise in what parameters were estimated in order for the model to converge. This was resolved pragmatically by fixing some parameters that were not central to the research questions or for which good quality information was available (Table 1).

**Table 1.** Fixed parameters used in the Bayesian hierarchical model

| Variable | Description | Parameter | JAGS code | Value | References |
| --- | --- | --- | --- | --- | --- |
| Birth | per capita birth rate | $b$ | b | 4 | [32, 39] |
| Survival | survival probability | $s$ | s | 0.908 | [32, 40] |
| Area (km<sup>2</sup>) | Size of region | $A$ | A | | |
| Shared border (km) | 15 x 15 matrix | $l$ | qb | | |
| Carrying Capacity | region 1 | $K_1$ | K[1] | 140000 | |
| | region 2 | $K_2$ | K[2] | 110000 | |
| | region 3 | $K_3$ | K[3] | 90000 | |
| | region 4 | $K_4$ | K[4] | 95000 | |
| | region 5 | $K_5$ | K[5] | 90000 | |
| Area (km<sup>2</sup>) | region 1 | $A_1$ | A[1] | 29479 | |
| | region 2 | $A_2$ | A[2] | 23180 | |
| | region 3 | $A_3$ | A[3] | 18146 | |
| | region 4 | $A_4$ | A[4] | 20446 | |
| | region 5 | $A_5$ | A[5] | 16172 | |

We were primarily interested in estimating four main parameters: the heterogeneity parameter for transmission *h*, the probability of an infected fox leaving an area *ρ*_*max*_, the observation rate *θ*, and the precision *τ* of the environmental noise (Table 2). The heterogeneity parameter *h* for transmission was given a gamma prior with mean 4 and variance 9. We experimented with the sensitivity of the model to the parameter *θ*, for quantifying case detection. We chose a prior for *ρ*_*max*_ (the rate at which infected foxes leave their focal region) where the posterior, informed by data, would have to move away from 0, ‘no movement’, to support connectivity between regions. Because infected foxes have a limited dispersal range [37], we expected the value of *ρ*_*max*_ to be small. To limit the search to a biologically plausible range of values for the leaving rate, we used a Beta prior with mean 0.004 and variance 0.000005 that declined with distance from 0. The precision of the environmental noise term *τ* was given a gamma prior with mean 1000 and variance 5000, based on fluctuations in fox reproductive effort reported in the literature [34]. This allowed the litter size (or birth pulse) to vary between 0.9 and 1.1 of the mean litter size each year.

**Table 2.** Priors used for the stochastic parameters in the Bayesian hierarchical model

| Variable | Parameter | JAGS code | Distribution | Prior | Mean and variance | Reference |
| --- | --- | --- | --- | --- | --- | --- |
| Leaving probability | $\rho$ | c | Beta | dbeta(1.26176, 156.45824) | (0.008, 5e-5) | |
| Heterogeneity parameter | $h$ | h | Gamma | dgamma(1.7777, 0.4444) | (4, 9) | |
| Vaccination Rate | $\nu$ | v | Beta | dbeta(12.3, 28.7) | (0.3, 0.005) | [35] |
| Environmental Noise | $\tau$ | tau | Gamma | dgamma(200, 0.2) | (1000, 5000) | [34] |
| Observation Probability | $\theta$ | obs1 | Beta | dbeta(9.45, 179.55) | (0.05, 0.00001) | |
| Population Region 1 |  | start_pop[1,1] | Gamma | dgamma(50.06, 0.0007) | (70750, 100e6) |  |
| Population Region 2 |  | start_pop[2,1] | Gamma | dgamma(30.95, 0.0006) | (55632, 100e6) |  |
| Population Region 3 |  | start_pop[3,1] | Gamma | dgamma(18.97, 0.0004) | (43550, 100e6) |  |
| Population Region 4 |  | start_pop[4,1] | Gamma | dgamma(24.08, 0.0005) | (49070, 100e6) |  |
| Population Region 5 |  | start_pop[5,1] | Gamma | dgamma(15.06, 0.0004) | (38813, 100e6) |  |

### 2 Model evaluation

To assess the model fit we used a probability integral transform, testing whether the observed rabies cases can reasonably be assumed to be arising from the chosen model. This was done by comparing the observed number of rabies cases to the posterior distribution of the expected number of infected cases estimated from the MCMC samples, calculating the percentile where the data point fell within the cumulative distribution function [38]. Because the credible intervals are calculated from the posterior of the expected values and not the posterior of the prediction, which also includes an error term, the credible intervals are more conservative and reflect only the uncertainty around the regression and not the prediction. As a result, they are narrower than the credible intervals of the prediction because they do not include this error.

To assess the model fit through time we plotted the computed monthly probability integral transform values through time. To explore the full range of potential rabies epidemic scenarios within the parameter space we simulated from the fitted model providing the same initial conditions including the size of the region, connectivity between regions, carrying capacity, initial number of susceptible and infected individuals, and rates of vaccination. We also assessed the model predictions when the model was provided with the first data point and the first 20 data points in each region using the probability integral transform to assess the prediction.

## Results

Across all regions the number of reported annual fox rabies cases ranged from 959 to 2375 pre-vaccination, with 2277 cases in 1990, the first year of vaccination. A swift decline in rabies followed, with 1643, 321, 28, 4 cases detected in the subsequent years. Fitting the model to the fox rabies case data generated key estimates of local transmission and spatial coupling between regions. The model captured key aspects of fox demography including the birth pulse, fluctuations in fecundity with environmental noise, and in surveillance, yielding a close fit to the case data in all 5 regions (Fig 1). From the fitted parameters, we estimated that less than 8% of the fox population were infected by rabies annually and that incidence was generally low. The model accommodated uncertainty in the biological and observation processes, allowing inference of missing time series of infected, susceptible, and vaccinated individuals by latent process methods.

**Fig 1.**
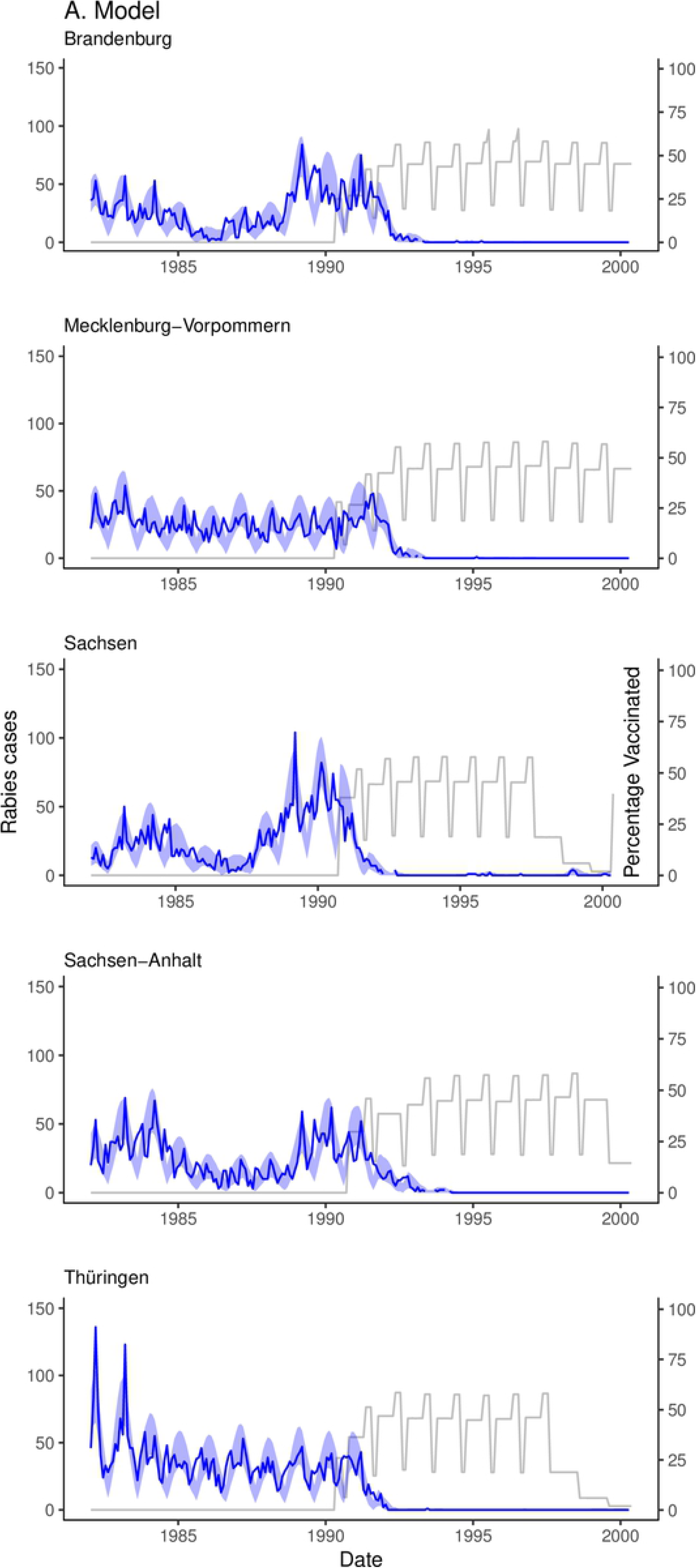
State-space model results for 5 federal states in Eastern Germany: Brandenburg, Mecklenburg-Vorpommern, Sachsen-Anhalt, Sachsen, and Thüringen. The gray line represents the estimated percentage of the population vaccinated. The dark blue line is the monthly reported rabies cases for each region. The light blue shaded region is the 95% credible intervals.

In terms of model fitting, the posterior for transmission heterogeneity *h* was both within the broad support of the prior and more precise than the prior, informed by the data (Fig 2). Too much flexibility in *θ* caused the model to explain most cases in terms of detectability, rather than epidemiological processes and tended to settle on posterior values that were too large compared to expert opinion for rabies and known limits of passive surveillance. Specifically, we found that wider priors for *θ* (e.g. mean = 0.10, variance = 0.01), went to biologically implausible areas of parameter space (around 30-40% of cases observed). The more flexible model also resulted in larger unrealistic values of *h* and poor model convergence, that we believe is due to the biological processes not being captured effectively. We addressed this pragmatically by incorporating expert opinion into our prior for detectability (mean = 0.05, variance = 0.00025).

**Fig 2.**
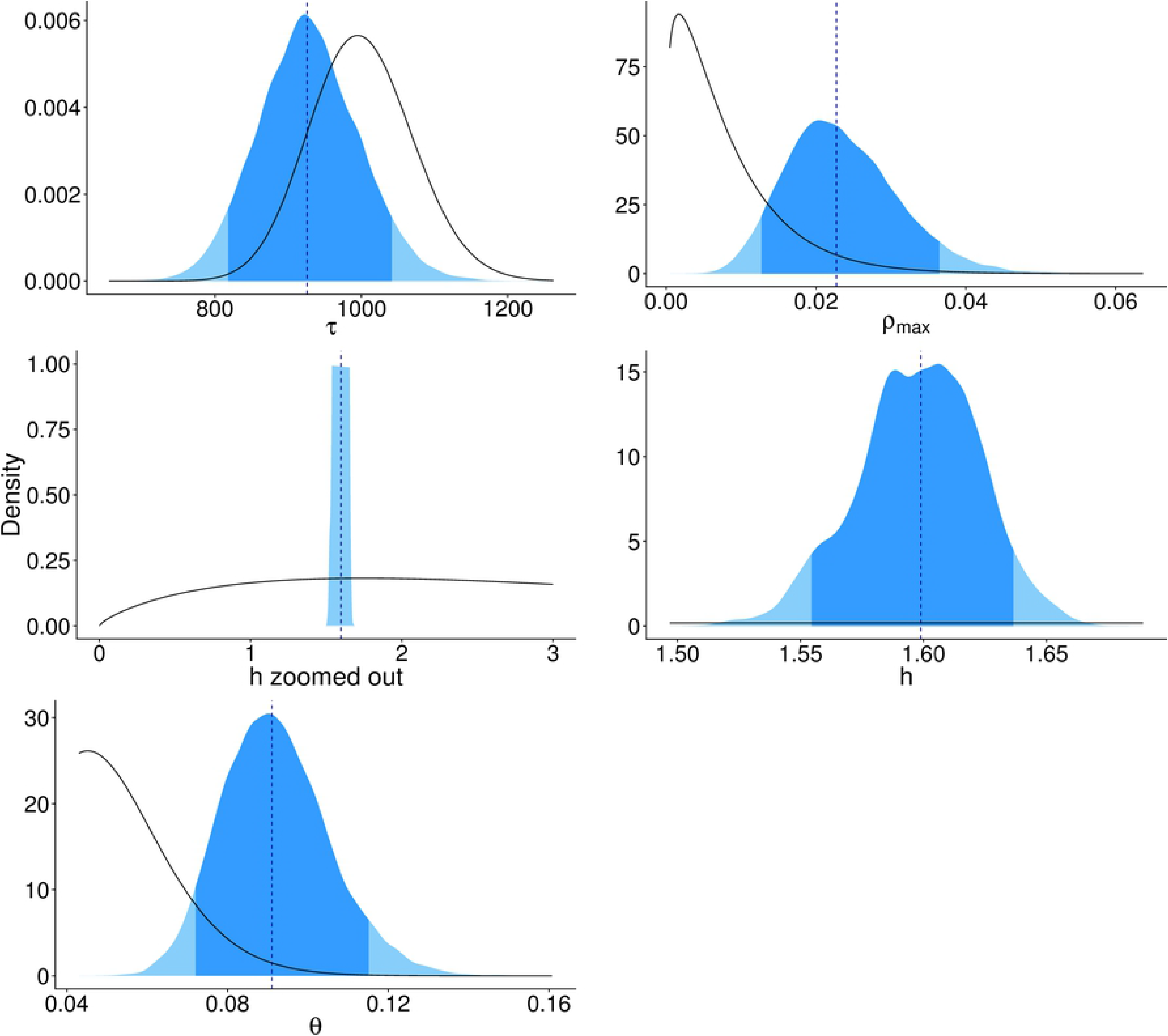
The posterior distribution and priors for the parameters estimated in the hierarchical Bayesian state-space model. The posterior distribution is shaded in blue (dark blue represents the 2.5-97.5% credible intervals, the light blue represents the 0-2.5% and 97.5-100% credible intervals), the black line represents the prior distribution. In order from top to bottom and left to right are the parameters representing fluctuations in fecundity due to environmental noise *τ*, rate of migration of rabid foxes between regions *ρ*_*max*_ in the equation, rabies transmission heteorgeneity *h*, zoomed out to show the prior, and zoomed in to show the distributions better, and the annual probability of observing rabies cases *θ*.

The posterior for *ρ*_*max*_ (the rate at which infected foxes leave their focal region) supported our hypothesis of connectivity between regions by moving away from 0. The posterior distribution of the environmental noise term *τ* shifts to the left of the prior, which suggests that the population fluctuates slightly more than specified by the prior. Although the fluctuations estimated from the model were larger than previously reported [34], they rarely exceeded 10% with the largest reaching 15%. This could be due to more extreme environmental changes during this period compared to when Lindström’s study was conducted.

The multi-year peaks in observed cases are largely explained by the annual birth rate and variation in both the observation rate and environmental noise (Fig 3). The environmental noise term influences the size of the susceptible population by increasing or decreasing the size of the birth pulse. In the model this results in more infected individuals, while a larger observation rate means that although more cases were detected, fewer cases occurred overall.

**Fig 3.**
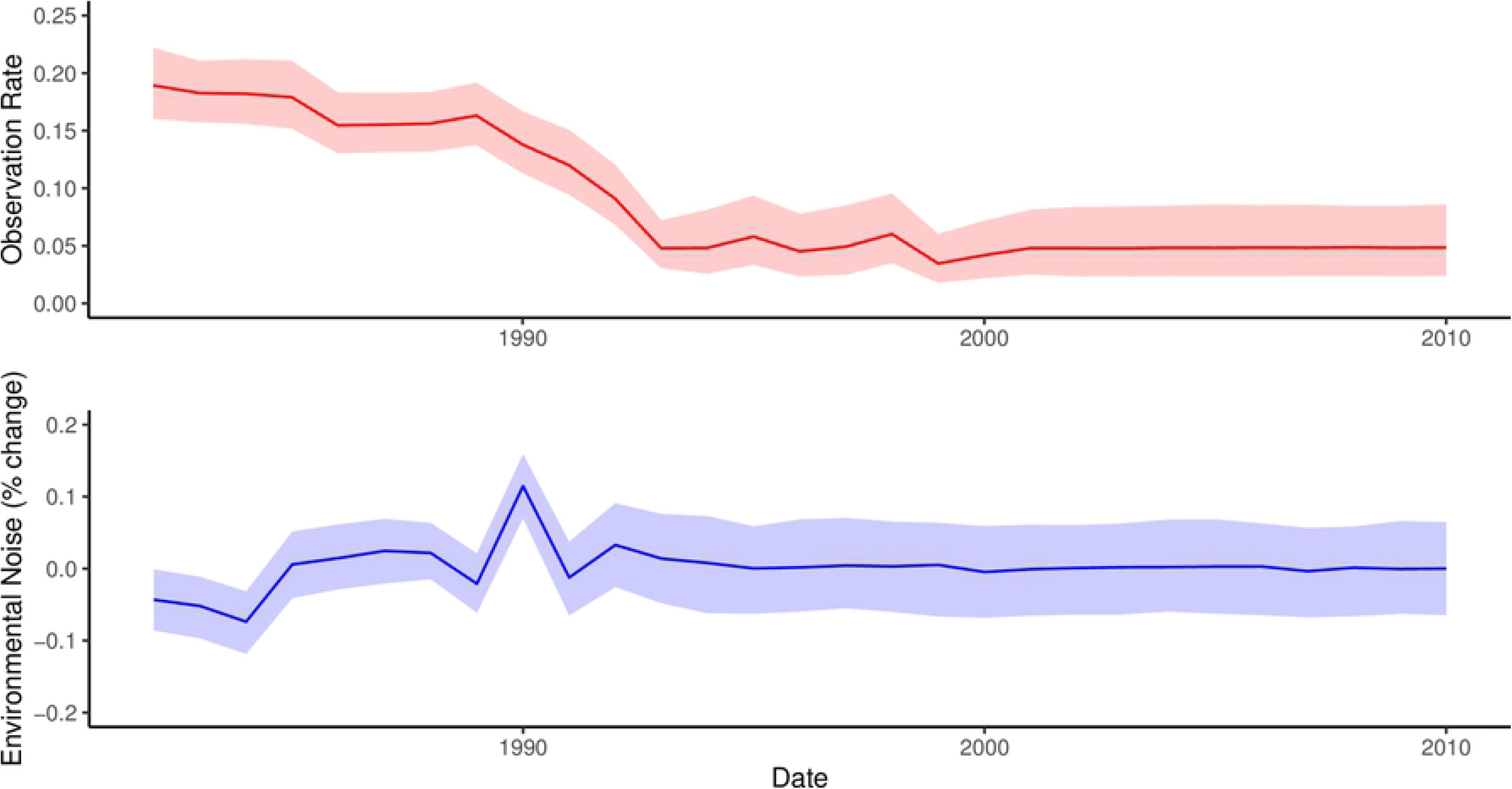
Annual observational and environmental noise estimates over time. The top and bottom panel show the annual observational noise estimates (red) and annual environmental noise estimates (blue) over time. The dark lines represent the mean estimates, the lighter envelope represents the 95% credible intervals. The environmental noise parameter fluctuates between 10 and +15%. The observational noise term ranges between 5% and 18%.

We estimated that incursions from neighbouring regions accounted for on average 1% or less of monthly rabies cases in a region, with the mean number of incursions varying between 0 and 4 per month in each region and from 0-24 per year (Fig 4. Low numbers of estimated external incursions from 2000 onwards reflect coincident declines in rabies cases in all 5 regions resulting from coordinated vaccination.

**Fig 4.**
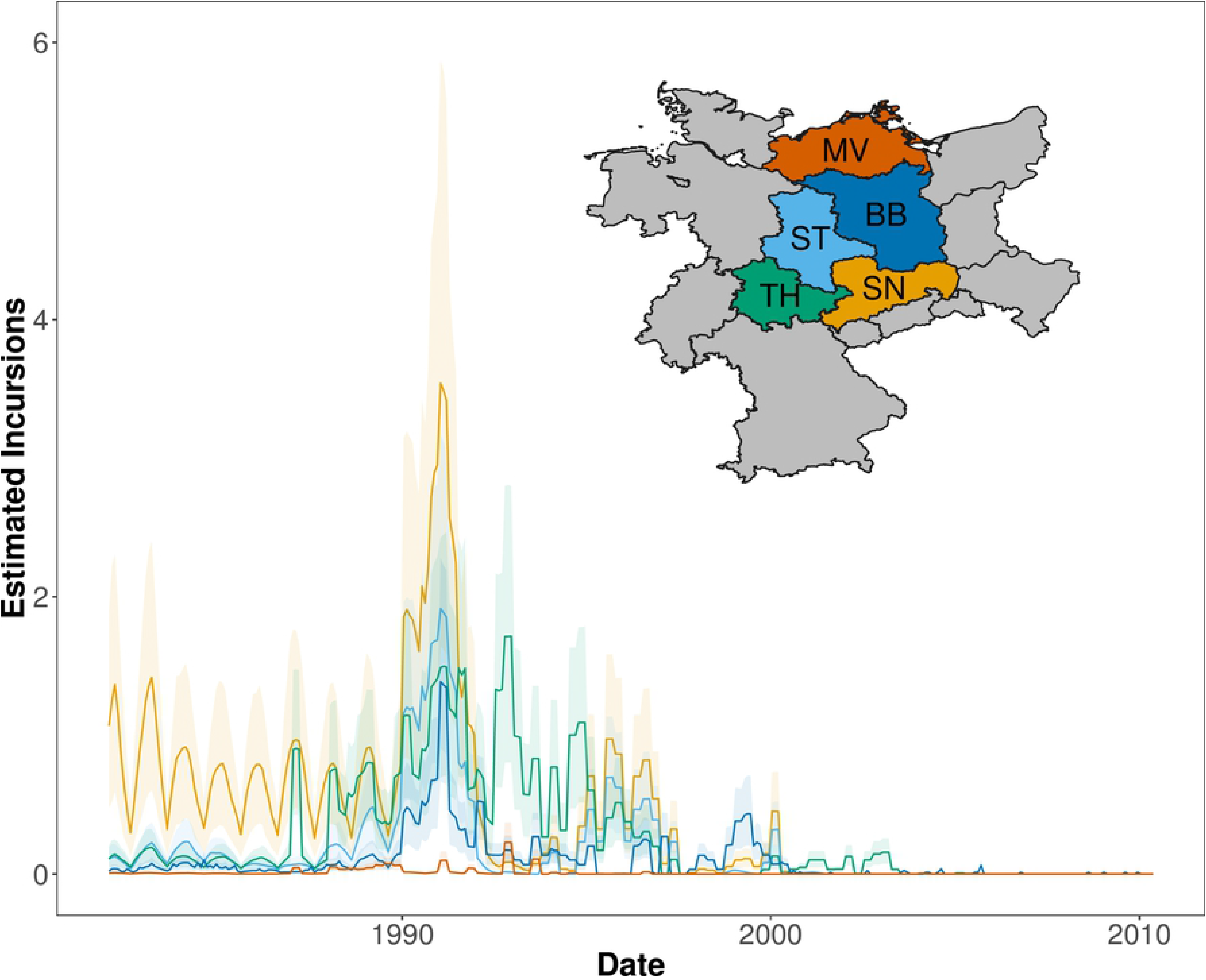
Estimated incursions per region through time. The map and incursions are colour-coded by region (BB = Brandenburg, MV = Mecklenburg-Vorpommern, SN = Sachsen, ST = Sachsen-Anhalt, and TH = Thüringen). Light grey regions represented neighbouring regions included in the model from Germany, Czech Republic, and Poland. The darker lines represent the mean number of incursions estimated per region. The lighter envelopes indicate the credible intervals.

Under the fixed vaccination rate (mean = 0.30, variance = 0.005), herd immunity peaked at between 60-75% of the population vaccinated in each region (Fig 1). Swift reductions in herd immunity occurred due to the entry of juvenile foxes into the population three months after the birth pulse (July), reducing the percentage of vaccinated individuals in the population by more than half. Between the annual entry of juvenile foxes, levels of herd immunity were maintained as vaccinated and susceptible individuals experience the same rate of natural mortality and rabies only claims a small proportion of the susceptible population (Fig 1).

From the plot of the probability integral transform calculated from the MCMC samples of the model fit, we can see that the model does not capture the largest peaks in observed cases and misses some of the observed cases between the birth pulses, when the population is at its lowest (Fig 5A). In the model, mortality is constant across the population, however, foxes experience a higher rate of mortality in the first months of life. Therefore, sharper population declines might be expected to follow the birth pulse due to this high juvenile mortality. Generally the fluctuations in observed cases were more variable compared to the model expectations. This could be because the model only allows for annual variability in the probability of detection and environmental variability rather than monthly variation. This was a deliberate modelling choice, as we found that too much flexibility in the detection probability or environmental noise (i.e. allowing these variables to vary by month or have a wider prior) resulted in overfitting and came at the expense of convergence of the other parameters, in particular transmission heterogeneity. Even though the fitted model expects a more gradual and smoothed number of cases than the observed data and the credible intervals are misleadingly narrow, the expectation is not far off from the observed cases and the model successfully captures the effect of vaccination.

**Fig 5.**
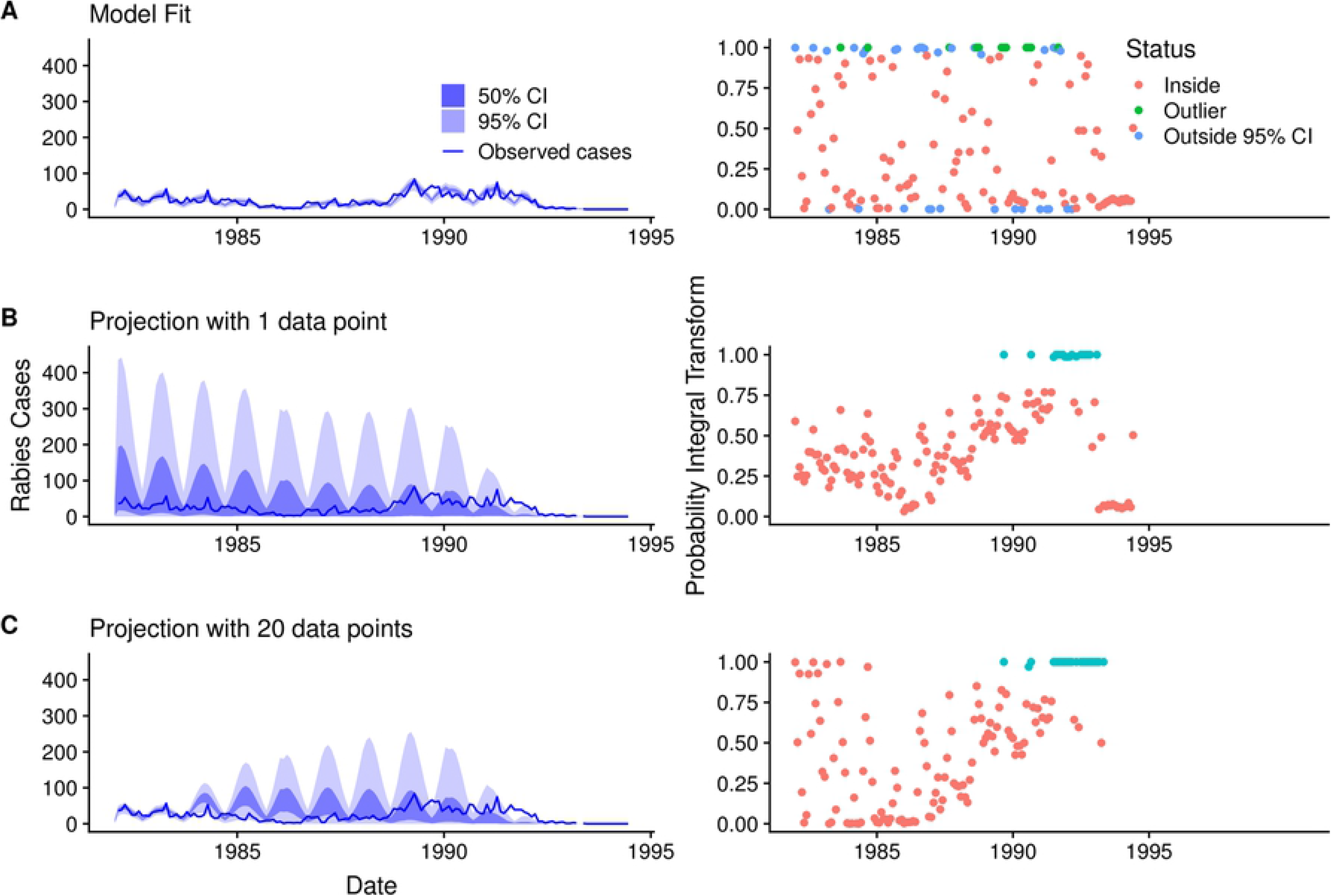
Model fit and projections with 1 and 20 data points with accompanying probability integral transform plots. The left panels show A. the model fit, B. the model projection with 1 data point, and C. the model projection with 20 data points. The right panels show the corresponding probability integral transform, i.e. where the observed case falls in the cumulative distribution function, over time for the three cases A-C. The dark blue line represents the observed rabies cases and the light and dark shaded blue regions represent the 95% and 50% Credible Intervals (CI) estimated from the MCMC samples, respectively. The colours in the probability integral transform plot indicate whether the observed cases falls inside (red) the 95% CI, outside (turquoise) the 95% CI but within the 100% CI, or outlier (green) when the case falls outside the CI entirely.

From the model projections, the most dominant features were oscillations due to the entry of juvenile foxes after the birth pulse (Figs 5B,C). In the first prediction, when the model is only provided with the first data point, all of the cases fall within the credible intervals and the majority of data points are within the middle 50% of the credible intervals, which suggests that the model prediction is too broad (Fig 5B). The model’s prediction improves when provided with the first 20 data points and is good at projecting the long-term behaviour and decline of rabies. However the prediction misses the tail end of the cases (Fig 5C). Although the credible intervals around the model predictions reduced when more data was provided, the model was unable to capture the multi-year oscillations in cases, which in the model fit were explained by annual fluctuations in environmental noise and variation in the annual observation rate (Fig 3).

## Discussion

Approaches to understand interactions between spatial and demographic processes are likely to reveal key insights into disease dynamics in wildlife populations. Here, we present a method to capture local transmission processes and spatial coupling between regions from partially observed data on wildlife diseases. Using a hierarchical Bayesian State-space metapopulation approach we were able to recreate observed dynamics, and infer missing time series by latent process methods. Specifically we find that (i) spatial coupling and local transmission can be estimated from aggregated data using a metapopulation modelling approach and a heterogeneous mixing term that captures the low incidence dynamics of rabies at a regional level; (ii) incursions of rabid foxes from other regions that account for less than 1% of cases are sufficient to trigger rabies re-emergence; and (iii) herd immunity achieved through bi-annual vaccination campaigns is short-lived due to fox population turnover. Together these findings have important practical implications for the design of control measures.

Partially observed disease data is common in the study of wildlife disease. Using a Bayesian state-space model we were able to accommodate uncertainty in the biological and observation processes and infer missing time series by latent process methods. By combining a metapopulation model representing space as a network of subpopulations with different rates of coupling, and a transmission process that approximates the scaling of individual interactions to regional dynamics, we provide a framework to estimate the rate of incursions and to model local dynamics. The heterogeneous mixing term involved a decay function that allowed for a reduction in transmission as the number of infected individuals increased. Our approximation was able to capture the low incidence of fox rabies and prevented unrealistically large epidemics. Earlier studies have accounted for heterogenous mixing in childhood diseases [17, 22] and influenza [18], however, this study is the first to use such an approximation for rabies. The approach has potential application for other diseases that circulate through local interactions but for which surveillance data is aggregated.

Incursions can play a crucial role in sustaining disease circulation, however many models do not explicitly consider between-region transmission. We estimated that incursions comprise less than 1 % of overall cases, reflecting the predominantly local nature of rabies transmission and the large size of the federal states. Although long-distance translocations of infected wildlife are known to occur [41], fox rabies transmission is thought to mainly result from movement of rabid and latently infected foxes. Even a small number of incursions from neighbouring regions through such movement can enable disease resurgence in a rabies-free region and are an important consideration in the design of vaccination strategies [42]. Coordinated vaccination efforts between regions can act to isolate foci of infection [43] and are crucial to the success of fox rabies elimination strategies [29]. Eastern Germany was able to rapidly eliminate rabies compared to elsewhere in Germany thanks to coordinated vaccination effort [29].

Herd immunity in foxes is dynamic as a result of demographic processes, declining by more than half following entry of juvenile foxes into the population following the birth pulse. Influxes of new susceptible individuals may result in isolated infections becoming rapidly reconnected with susceptible hosts and transmission maintained in the absence of adequate immunity [Katie- do you have a suggestion for a citation]. Our analyses of vaccination suggest that herd immunity is only maintained through regular ORV campaigns due to high population turnover. This may also have important implications for other wildlife diseases with marked birth pulses [44].

While our model generally captured rabies dynamics, it struggled to capture monthly variation in cases and long-term multi-year dynamics. The strongest mechanistic feature of our model driving these fluctuations was the entry of juvenile foxes into the population after the birth pulse. This is evident from both model projections with 1 or 20 data points (Figs 5B,C). Although the credible intervals around the model predictions reduced when more data was provided (20 data points compared to 1 data point), the model did not able to capture multi-year oscillations in cases, which were explained by the annual environmental and observation noise parameters. It is probable that changes in resource availability and variation in detection explains some of the variation, but, we did not have any data to inform these parameters aside from expert opinion. We speculate that nuances in the dynamics may be obscured due to the aggregation of data. The behaviour of the observation parameter *θ* over time suggests either marked changes in case detection or other mechanisms not being captured that the model is apportioning to the observation rate. In analysing the data at a finer spatial scale we observe multiple foci of cases within a region. However, because the data are aggregated these local dynamics are not evident. Nonetheless, the model expectation is not far off from the observed cases and the model successfully captures the effect of vaccination. Therefore, although the model fit is overconfident, overall it does a good job of capturing the disease dynamics.

## Conclusion

Disease dynamics play out across space, irrespective of borders. Modelling approaches to understand interactions between spatial and demographic processes can reveal key insights into disease dynamics and are crucial to the planning of regional vaccination strategies. Several studies have been central for guiding control of fox rabies in Europe. A simple deterministic, compartmental model based on fox population biology compared the dynamics of rabies under culling versus vaccination [39]. Spatially explicit individual-based models (IBM) of rabies have since been used to evaluate vaccination strategies for elimination [40, 45] and emergency vaccination under limited resources [46, 47]. Our model adds to this body of work and is the first to estimate epidemiological parameters from fitting to rabies incidence data. Our study adds to the considerable body of work that has been central for guiding the control of fox rabies in Europe [39, 46, 47]. It is the first to estimate key epidemiological parameters, including spatial coupling and local transmission, using a model fit to data. Our findings have implications for strategies aiming to achieve and maintain rabies freedom and the modeling approach can be used to further explore vaccination strategies to inform ongoing vaccination in Eastern Europe [42]. This work also makes an important methodological contribution to the study of spatial disease dynamics in other wildlife diseases where only limited epidemiological and demographic data are available.

## Supporting information

**S1 Fig. Model Diagnostics** Posterior and prior distributions and traceplots for all parameters in the model.

## Acknowledgments

We gratefully acknowledge the technical assistance provided by Ronald Schröder and Patrick Wysocki from the Friedrich Loeffler Institute in accessing the data.

